# Secretin release after Roux-en-Y Gastric Bypass reveals a population of glucose-sensitive S-cells in distal small intestine

**DOI:** 10.1101/791335

**Authors:** Ida M. Modvig, Daniel B. Andersen, Kaare V. Grunddal, Rune E. Kuhre, Christoffer Martinussen, Charlotte B. Christiansen, Cathrine Ørskov, Pierre Larraufie, Richard Kay, Frank Reimann, Fiona M. Gribble, Bolette Hartmann, Kirstine N. Bojsen-Møller, Sten Madsbad, Nicolai J. Wewer Albrechtsen, Jens J. Holst

## Abstract

**Objective:** Gastrointestinal hormones contribute to the beneficial effects of Roux-en-Y gastric bypass surgery (RYGB) on glycemic control. Secretin is secreted from duodenal S-cells in response to low luminal pH, but it is unknown whether its secretion is altered after RYGB and if secretin contributes to the post-operative improvement in glycemic control. We hypothesized that secretin secretion increases after RYGB as a result of the diversion of nutrients to more distal parts of the small intestine, and thereby affects islet hormone release.

**Methods:** A specific secretin radioimmunoassay was developed, evaluated biochemically, and used to quantify plasma concentrations of secretin in 13 obese individuals before, 1 week after and 3 months after RYGB. Distribution of secretin and its receptor was assessed by RNA-sequencing, mass-spectrometry and *in situ* hybridization in human and rat tissues. Isolated, perfused rat intestine and pancreas were used to explore the molecular mechanism underlying glucose-induced secretin secretion and to study direct effects of secretin on glucagon, insulin and somatostatin secretion. Secretin was administered alone or in combination with GLP-1 to non-sedated rats to evaluate effects on glucose regulation.

**Results:** Plasma postprandial secretin was more than doubled in humans after RYGB (P<0.001). The distal small intestine harbored secretin expressing cells in both rats and humans. Glucose increased secretion of secretin in a sodium-glucose co-transporter dependent manner when administered to the distal part but not into the proximal part of the rat small intestine. Secretin stimulated somatostatin secretion (fold change: 1.59, P<0.05) from the perfused rat pancreas but affected neither insulin (P=0.2) nor glucagon (P=0.97) secretion. When administered to rats *in vivo*, insulin secretion was attenuated and glucagon secretion increased (P=0.04), while blood glucose peak time was delayed (from 15 min to 45 min) and gastric emptying time prolonged (P=0.004).

**Conclusion:** Glucose-sensing secretin cells located in the distal part of the small intestine may contribute to increased plasma concentrations observed after RYGB. The metabolic role of the distal S-cells warrants further studies.

## 1. Introduction

In the duodenum, enteroendocrine S-cells secrete secretin in response to luminal acidification and thereby regulate the intraluminal milieu by stimulating the exocrine pancreas to secrete water and bicarbonate (1). Secretin, however, also has other actions including inhibition of gastric emptying (2–4) and has been suggested to regulate secretion of the pancreatic islet hormones (5–10).

Gastrointestinal hormones, including glucagon-like peptide-1 (GLP-1), are currently used for treatment of metabolic diseases and endogenous GLP-1 contributes to the glucose lowering and weight reducing effect of Roux-en-Y gastric bypass (RYGB) (11,12). Whereas it is well established that GLP-1 secretion is increased after RYGB in humans (13), it is unknown if secretin secretion is increased.

Glucose is a powerful stimulus for GLP-1 secretion (14) and is rapidly absorbed in the upper part of the small intestine, therefore intraluminal concentrations are low in the distal part of the small intestine. We hypothesized that delivery of glucose to the distal small intestine, as seen after the diversion following RYGB, stimulates secretin secretion by luminal-sensing mechanisms. To address this, we measured plasma secretin before and after RYGB. Furthermore, we isolated and perfused the upper or lower half of the rat small intestine and studied the effect of intra-luminal glucose on secretin secretion as well as the molecular sensors involved.

To investigate potential effects of secretin on blood glucose and secretion of islet hormones, we administered secretin subcutaneously to conscious rats and perfused, in separate experiments, the rat pancreas, testing the effects of secretin on glucagon, insulin and somatostatin secretion.

## 2. Materials and Methods

### 2.1. Ethical Approvals

#### Human studies

Written informed consent was obtained from all study participants, and the study was approved by the Municipal Ethics Committee of Copenhagen in accordance with the Helsinki-II declaration and by the Danish Data Protection Agency, and registered at www.clinicaltrials.gov (NCT01993511).

#### Animal Studies

Animal studies were conducted with permission from the Danish Animal Experiments Inspectorate, Ministry of Environment and Food of Denmark, permit 2018-15-0201-01397, and in accordance with the EU Directive 2010/63/EU and guidelines of Danish legislation governing animal experimentation (1987), the National Institutes of Health (publication number 85-23), and approved by the local ethical committee (EMED, P18-336).

### 2.2. Peptides

GLP-1 7-36NH_2_, rat and human secretin were obtained from Bachem (cat no. 4030663, cat no. 4037181 and cat no. 4031250, Bubendorf, Switzerland). Radioactive labeled rat and human secretin was obtained from Phoenix Pharmaceuticals, Inc (cat no. T-067-06 and T-067-07, CA, USA). Development and evaluation of a secretion radioimmunoassay are described in Supplementary Materials 1.

### 2.3. Mixed meal tests before and after Roux-en-Y Gastric Bypass surgery in obese subjects

Plasma obtained during a standardized liquid mixed-meal test (MMT) before, one week and three months after RYGB from 13 obese subjects (type 2 diabetes; n=4, impaired glucose tolerance; n=3, normal glucose tolerance; n=6) was analyzed. Glucose tolerance was determined by standard OGTT preoperatively. All samples were from a study by Martinussen et al as previously described (15).

### 2.4. Distribution of secretin and GLP-1 along the gastrointestinal tract in humans and rats

Mass-spectrometry based detection was used to assess distribution of secretin and GLP-1 (for comparison) in human intestinal tissue as described previously (16). The secretin and GLP-1 profiles are presented as peak area divided by tissue weight.

Whole wall tissue biopsies (~1 cm) of esophagus, ventricle, duodenum, proximal jejunum, distal ileum and colon were collected from non-fasted rats (anatomical definitions are listed in Supplementary Table 1). Peptides in tissue biopsies were extracted using trifluoroacetic acid (Supplementary Materials 2), and immunohistochemical staining of secretin-positive cells was performed on paraffin embedded tissue samples using anti-secretin (5585-3), a generous gift from Professor Jan Fahrenkrug (Supplementary Materials 3).

### 2.5. Animal experiments

Male Wistar rats (200-250 g) were obtained from Janvier (Le Genest-Saint-Isle, France) and housed two to four rats per cage. Rats were allowed one week of acclimatization and kept on a 12 h light/dark cycle with ad libitum access to water and standard chow.

#### 2.5.1 Isolated perfused rat small intestine and pancreas

Perfusion was performed using a single pass perfusion system (UP100, Hugo Sachs Harvard Apparatus, Germany). The rat small intestine and pancreas were surgically isolated (as described in Supplementary Materials 4). Each protocol started with a baseline period followed by addition of various test substances. Test stimulants used in the perfused intestine included 0.1 M HCl, 0.1 M NaHCO_3_^−^, 0.05 M KCl (Sigma-Aldrich, Brøndby, Denmark), 20% (w/v) D-glucose (cat no. G8270, Sigma-Aldrich, Denmark) and 10 mmol/L Phloridzin (cat no. P3449, Sigma-Aldrich, Denmark). Test stimulants used in the perfused pancreas were 1 nmol/L GLP-1 7-36NH_2_ and both 20 pmol/L and 2 nmol/L secretin.

#### 2.5.2 In vivo experiments in rats

Experiments were carried out on 2 occasions on fasted rats (300±13 g) just before their nocturnal feeding period (5:00 PM). Rats were divided into weight-matched groups (n=8/group). At −10 minutes, 200 μL tail blood was collected into pre-chilled EDTA-coated capillary tubes (catalog no. 200 K3E, Microvette; Sarstedt, Nümbrecht, Germany) and instantly transferred onto ice. At −5 minutes, 300 uL test solution was injected subcutaneously. Test solutions were isotonic saline, (negative control) or peptides prepared in isotonic saline: secretin (30 nmol/kg), GLP-1 (30 nmol/kg) or secretin+GLP-1 (both 30 nmol/kg). At 0 minutes, a bolus of D-glucose (2 g/kg) and acetaminophen (100 mg/kg), prepared in isotonic saline, was given orally. Rats from the same cage received different treatments. Blood was collected at times −10, −5, 5, 15, 30, 45, 60 and 90 min. Glucose was measured immediately after blood collection, while the remainder of the samples were instantly transferred onto ice and centrifuged (1,650 g, 4°C, 10 min) within half an hour to obtain plasma. Plasma was transferred to Eppendorf tubes, immediately frozen on dry ice and stored at −20°C until analysis.

#### 2.5.3 Biochemical measurements of perfusion effluents, rat blood and rat plasma

Perfusion effluents: Insulin was measured with antiserum (code no. 2006-3) which cross-reacts with rodent insulin I and II (17). Glucagon was measured with an antibody directed against the C-terminus (code no. 4305) as previously described (18). Total GLP-1 (the sum of 7-36NH_2_, 9-36NH_2_ and potential mid-terminal cleaved fragments) was measured using a C-terminal specific radioimmunoassay targeting amidated forms (code no. 89390) (19). Somatostatin was measured using a side-viewing antibody (code no. 1758-5), detecting all bioactive forms of somatostatin (20,21).

Blood glucose was measured using a glucometer (Accu-Chek Mobile, catalog no. 05874149001; Roche Diagnostics, Mannheim, Germany). Plasma concentrations of glucagon were measured by sandwich ELISA (catalog no. 10-1281-01; Mercodia AB, Uppsala, Sweden) (22). Plasma concentrations of insulin and C-peptide were measured using ELISAs (catalog no. 10-1250-01 and 10-1172-01; Mercodia AB, Uppsala, Sweden).

### 2.6. RNA sequencing of human islets

Publicly available RNAseq dataset (from GSE85241, GSE81608 and E-MTAB-5061 (23–25)) were obtained. Average Reads Per Kilobase Million (RPKM) values for secretin receptor (SCTR) and GLP-1 receptor (GLP-1R) were uploaded to the Jupyter Notebook (http://jupyter.org/). Data were then log2 transformed and mean expression levels were calculated in alpha, beta and delta cells, respectively. We excluded individuals with diabetes from these analyses. For further details about the donors, isolation of cells and RNA-sequencing methods please see the original studies (23–25).

*In situ* hybridization on rat pancreases is described in Supplementary Materials 5.

### 2.7. Statistical analysis

Clinical samples and in vivo data: Plasma concentrations of hormones were evaluated using area under the curve analysis and statistical testing by one-way ANOVA followed by Holm-Sidak multiple comparisons test. A mixed-effect model was applied in order to test the effect of treatment and time on the variables.

Perfusion experiments: Data are expressed as mean±SEM concentrations in venous effluents (pmol/L). Since the perfusion flow was kept constant through the experiments, this mirrors the actual secretion output (which can be calculated by multiplying with the flow rate). To test for statistical significance, mean values within the test period (based on 10 subsequent minutes) were compared with mean values in the baseline period (10 subsequent minutes prior to test stimulant administration) using statistical testing as described above.

Calculations were made using STAT14 (SE), College Station, Texas, USA. For illustrations, GraphPad Prism version 8.0 (GraphPad Software, La Jolla California USA) and Adobe CC software suite were used (San Francisco, CA, USA). P-values < 0.05 were considered statistically significant. Data are shown as means±SEM.

## 3. Results

### 3.1. Development of a sensitive secretin assay

We generated titer curves for three different antibodies against human secretin (structure in Figure 1A) using two radioactive iodine-labeled secretin peptides (termed tracer). Based on binding characteristics, the antibody named 5595-3 was selected for further testing and for preparation of calibrator curves (Supplementary Figure 1). By sequencing alignment, we found that human and rat secretin differ at position 14-16 (human: REG vs. rat: QDS) (Figure 1A) but no species variation was found within the antibody’s epitope (position 18-27) (Figure 1A). For rat samples, we therefore used the same antibody and tracer. For calibrating purposes, we included the rat isoform of secretin as calibrator control.

**Figure 1:**
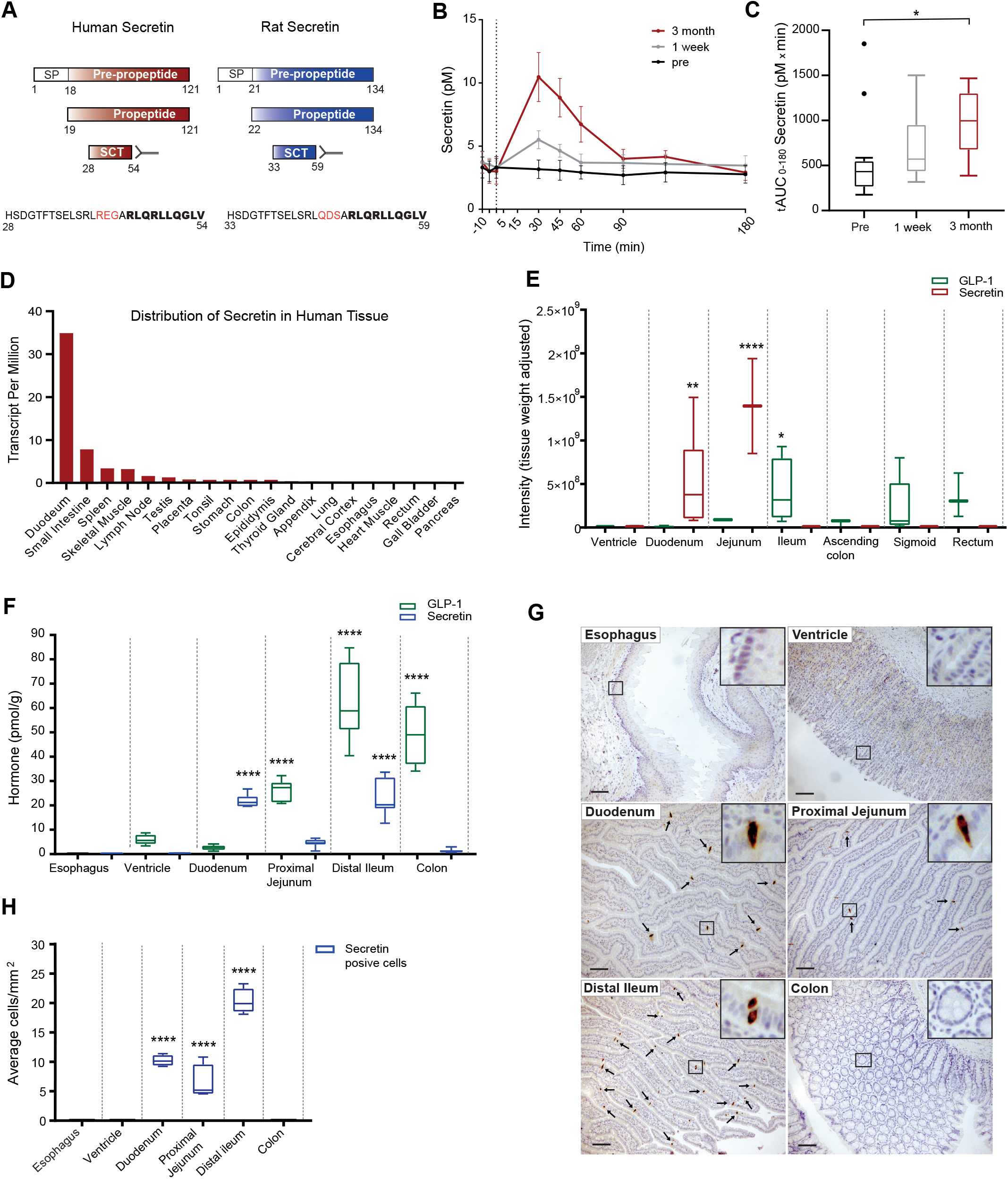
Secretion of secretin after Roux-en Y Gastric Bypass surgery and distribution of secretin in rats and in humans. **A:** Structure of human and rat secretin and antibody binding in the secretin assay. SP: signal peptide. Red capitals: differences between rat and human peptide sequences. Highlighted capitals: Ab epitope. **B:** Postprandial plasma concentrations of secretin before (pre), one week (1 week) and three months (3 month) after RYGB surgery in obese individuals, n=13. **C:** Calculated total area under the curve (tAUC_0-180_) is shown for panel B. **D:** Expression profiles of secretin in human tissues. RNA-seq data from human tissues presented as Transcripts Per Million (TPM) (26). **E:** Distribution of secretin and GLP-1 in human intestinal tissue. Mass-spectrometry profiling of secretin and GLP-1 in the human gastrointestinal tract. Levels of secretin and GLP-1 are presented as peak area divided by tissue weight, n=2-9 (16). **F:** Extractable secretin and GLP-1 concentrations in whole-wall biopsies along the rat gastrointestinal tract (pmol/g tissue), n=7 male Wistar rats. **G:** Immunohistochemical staining of secretin in the rat gastrointestinal tract (Ab:5585-3, 1:20.000), n=4 male Wistar rats. **H:** Counting of secretin-positive cells (immunohistochemistry) along the rat gastrointestinal tract (cells/mm^2^), scale bar 100 μm, n=4 male Wistar rats. Data are shown as means ± SEM. P>0.05. *P<0.05, **P<0.01, ****P<0.0001 by a One-way ANOVA with correction for multiple testing (Holm-Sidak).

Recovery of human secretin added to human pooled plasma was calculated to 71±11% (mean±SD) with a lower limit of detection of 1 pmol/L and a dynamic range from 2.5 pmol/L to 80 pmol/L (Supplementary Figure 1) using solvent phase-extraction (70% ethanol). We did not observe cross-reactivity (not significantly different compared to background) towards CCK, glucagon, glicentin, GLP-1, human insulin, oxyntomodulin, neurotensin and PYY at concentrations up to 300 pmol/L.

#### 3.1.1. Comparison of Commercial ELISAs

The two commercially available ELISAs had low recoveries of human secretin of 15%±8% (mean±SD) in assay buffer. A Bland-Altman analysis (comparing the concentrations measured using the two ELISAs to the in-house developed RIA) showed acceptable (less than 2SD) degree of variation for concentrations above 20 pmol/L, however, for physiological plasma concentrations of secretin (<20pmol/L) the two commercial ELISAs were inadequate (Supplementary Figure 2).

Having developed a specific and sensitive secretin radioimmunoassay, we next investigated whether postprandial plasma secretin concentrations are elevated after RYGB surgery.

### 3.2. Roux-en-Y Gastric Bypass surgery increases meal-induced secretin release in obese individuals

Fasting plasma concentrations of secretin were not significantly different after comparison with before RYGB (Figure 1B, P>0.45), whereas postprandial plasma secretin concentrations in response to a liquid meal test were significantly increased three months after compared with before RYGB (P<0.05), reaching a 2-3 fold higher peak value, and total AUCs were approximately doubled (tAUC_0-180min_: 569±475 vs. 956±360 pmol/L x min) (Figure 1C). Concentrations were also increased one week after RYGB but to a more modest degree and total AUCs did not differ from pre-operative AUCs (P=0.67).

### 3.3. Distribution of secretin in humans and rats

Using publicly available RNA-seq data from 37 different human tissues (26) expression profiles of secretin showed a maximum in the duodenum, as expected. However, high levels were also found in the more distal parts of the small intestine (Figure 1D), consistent with previous reports on secretin distribution in other species (27,28). Mass-spectrometry analysis on human gastrointestinal tissue revealed that secretin concentrations were highest in the duodenum as well as in jejunal biopsies (Figure 1E). Concentrations of GLP-1 7-36NH_2_ were included as positive controls. GLP-1 concentration was highest in the distal part of the gastrointestinal tract (ileum, colon and rectum) (Figure 1E) in line with previous reports (29).

To examine if the distribution of secretin in rats was similar to humans, we measured the concentration of extractable secretin from esophagus to colon in rats (Supplementary Table 1). Secretin was not detectable in esophagus and the ventricle (Figure 1F, *n=7*) but in the duodenum and in the distal ileum, secretin was found at comparable concentrations ~20-30 pmol/g tissue (Figure 1F, *n=7*). Concentrations of GLP-1 increased from duodenum to colon with the highest concentration in the distal ileum (~60 pmol/g tissue) (Figure 1F, *n= 7*) in line with a previous report (30). Immunohistochemical stainings of secretin (Figure 1G+H) showed no staining in esophagus, ventricle and colon and high intensity in proximal jejunum and distal ileum, consistent with the measured extractable concentrations.

### 3.4. Secretin responses from the proximal and distal small intestine using the isolated perfused rat intestine model

Luminal infusion of HCl increased secretin secretion 4-fold (Baseline: 3.7±0.9 vs HCl: 15.6±2.3 pmol/L, P<0.05, *n=6*) (Figure 2A+B). Following HCl infusion, the intestine was flushed with sodium bicarbonate (0.1 M) to neutralize luminal pH (Figure 2A). This returned secretin secretion to pre-HCl-stimulatory levels. KCl (0.05 M) was infused intra-vascularly at the end of the experiment (Figure 2A). Secretion of secretin increased significantly compared to preceding baseline secretion (Baseline: 5.2±0.8 vs. KCl: 9.1±0.9 pmol/L, P<0.05, *n=6*) (Figure 2B). Our data therefore suggests that cell depolarization is involved in secretin release from the perfused rat small intestine and furthermore that our perfusion model reflects the physiology of secretin secretion. We therefore decided to evaluate potential differences in HCl and glucose-induced secretin secretion in the proximal and distal small intestine using this model.

**Figure 2:**
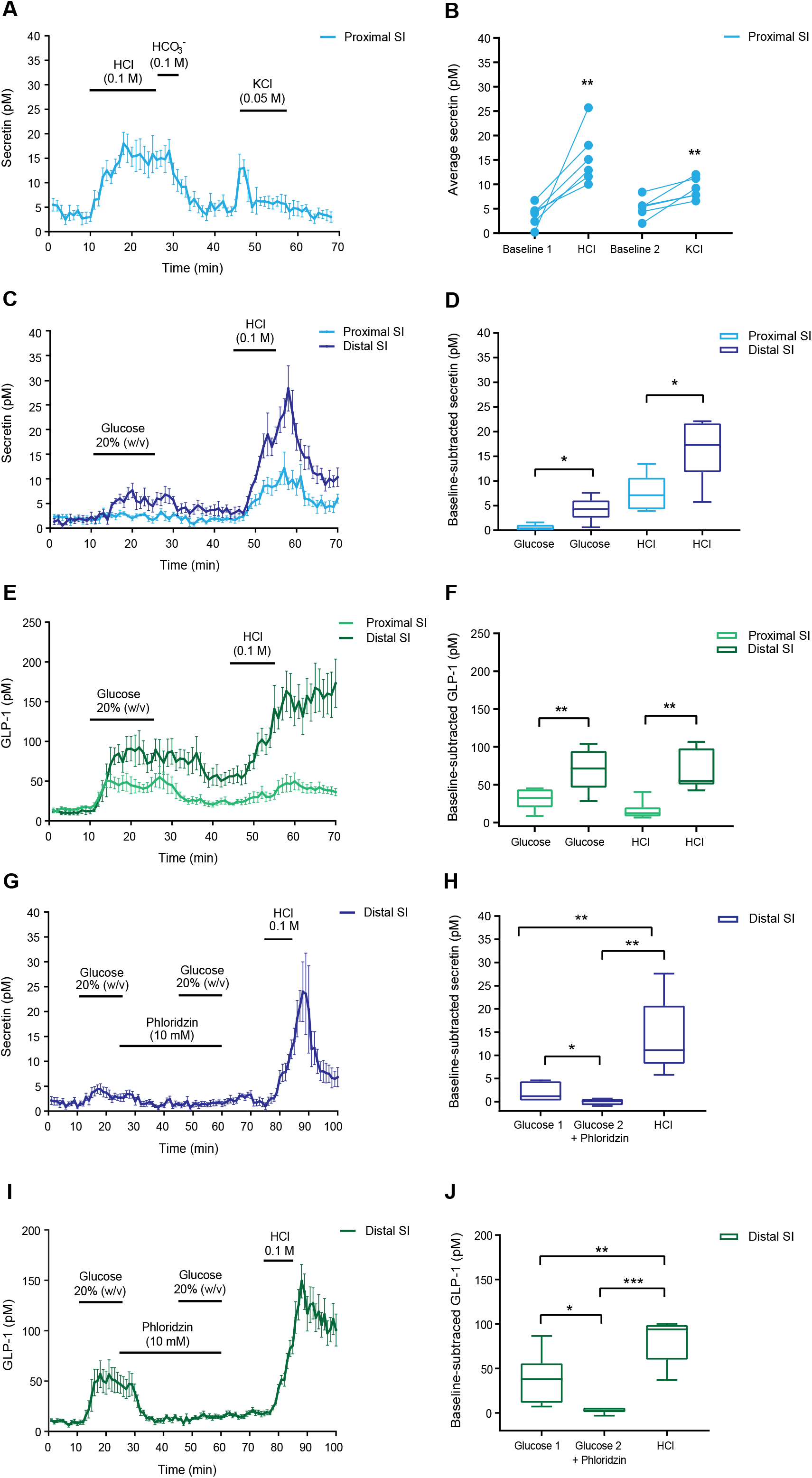
Differential sensing of glucose by secretin secreting cells in the perfused proximal vs distal rat small intestine. **A-B: Validation of the perfused rat intestine as a model to study secretin release.** HCl (0.1 M) was infused intra-luminally between minute 11-25, NaHCO_3_^−^ (0.1 M) was infused between minute 26-30. KCl (0.05 M) was infused intra-vascularly between minute 46-60. **A:** Secretin concentrations in venous effluents (pmol/L). **B:** Mean concentrations of secretin at baseline (Baseline 1 and 2) and following stimulation with either luminal HCl or vascular KCl. **C-F: Differential sensing of glucose in the proximal vs. distal intestine.** Glucose (20% w/v) was intra-luminally infused between minute 11-25 in either the proximal or the distal isolated perfused rat small intestine. HCl (0.1 M) was intra-luminally infused between minute 46-55 as a positive control for secretin release. **C:** Secretin concentrations (pmol/L). **D:** Mean baseline-subtracted secretin concentrations in response to glucose and HCl stimulation. **E:** GLP-1 concentrations (pmol/L). **F:** Mean baseline-subtracted GLP-1 concentrations in response to glucose and HCl stimulation. **G-J.: Sodium-coupled glucose transporters (SGLT) are involved in distal glucose sensing.** Glucose (20% w/v) was infused intra-luminally between minute 11-25 and between minute 46-60. Phloridzin (10 mmol/L), an SGLT inhibitor, was infused intra-luminally between minute 26-60. HCl (0.1 M) was infused intra-luminally between minute 76-85 as a positive control for secretin release. **G:** Secretin concentrations (pmol/L). **H:** Mean baseline-subtracted secretin concentrations in response to HCl and glucose stimulation with and without phloridzin. **I:** GLP-1 concentrations (pmol/L). **J** : Mean baseline-subtracted GLP-1 concentrations in response to HCl stimulation and glucose stimulation with and without phloridzin. Data are shown as means ± SEM. P>0.05. *P<0.05, **P<0.01, ***P<0.001, by a One-way ANOVA with correction for multiple testing (Holm-Sidak), n=6 male Wistar rats for all groups.

#### 3.4.1. Differential sensing of glucose by secretin producing cells in the proximal and distal small intestine

Luminal HCl infusion increased secretin secretion (P<0.05) from both the proximal and distal half of the small intestine, but to the greatest extent in the distal half (baseline-subtracted values; Proximal. HCl: 7.6±1.5 vs. Distal. HCl: 16.3±2.6 pmol/L, P<0.05, *n=6*) (Figure 2C+D). Of particular notice, distal, *but not* proximal, infusion of glucose resulted in a significant (P<0.05) secretin response (baseline-subtracted values; Proximal. Glucose: 0.48±0.3 vs Distal. Glucose: 4.3±1 pmol/L, *n=6*) (Figure 2C+D) suggesting that distally located secretin cells are capable of sensing glucose.

GLP-1 secretion was measured as this is a well validated control in this experimental setup (31,32). Secretion of GLP-1 in response to both glucose and HCl infusion was likewise significantly higher from the distal small intestine compared to the proximal small intestine (Figure 2E+F), which is consistent with high extractable concentrations of GLP-1 in the distal small intestine (Figure 1F).

#### 3.4.2. Glucose induced secretin secretion from the distal small intestine is mediated by sodium-glucose co-transporter

Inhibition of SGLT with the competitive SGLT1/2 inhibitor, phloridzin, eliminated glucose-stimulated secretin response from the distal rat small intestine (baseline-subtracted values; Glucose: 1.9±0.7 vs. Glucose + Phloridzin: 0±0.2 pmol/L, P<0.05, *n=6*) (Figure 2G+H), indicating that sodium-coupled glucose absorption is involved in the mechanism of glucose stimulated secretin secretion from the distal rat small intestine.

Consistent with previous reports (33,34), phloridzin eliminated glucose-stimulated GLP-1 secretion from the distal rat small intestine (baseline-subtracted values; Glucose: 38.5±10.4 vs. Glucose+Phloridzin: 2.6±1.2 pmol/L, P<0.05, *n=6*) (Figure 2I+J). HCl was included as a positive control for secretin secretion in the end of each experiment.

### 3.5. Secretin delays blood glucose peak time and increases plasma glucagon in non-sedated rats

Blood glucose concentrations were similar across groups at baseline (P>0.71). Peak times for blood glucose were prolonged from 15 min (saline group) to 30 min (GLP-1), 45 min (secretin), and 60 min (secretin+GLP-1) (Figure 3A) but the incremental area under the curve (iAUC_0-90min_) was not significantly different between groups (P>0.77, Figure 3B). Gastric emptying, reflected by changes in plasma acetaminophen concentrations, was markedly prolonged in secretin treated rats compared to saline and GLP-1 treated groups (P<0.05, Figure 3C+D). GLP-1 had significant effect on plasma acetaminophen at the initial time points compared to saline (P<0.05).

**Figure 3.**
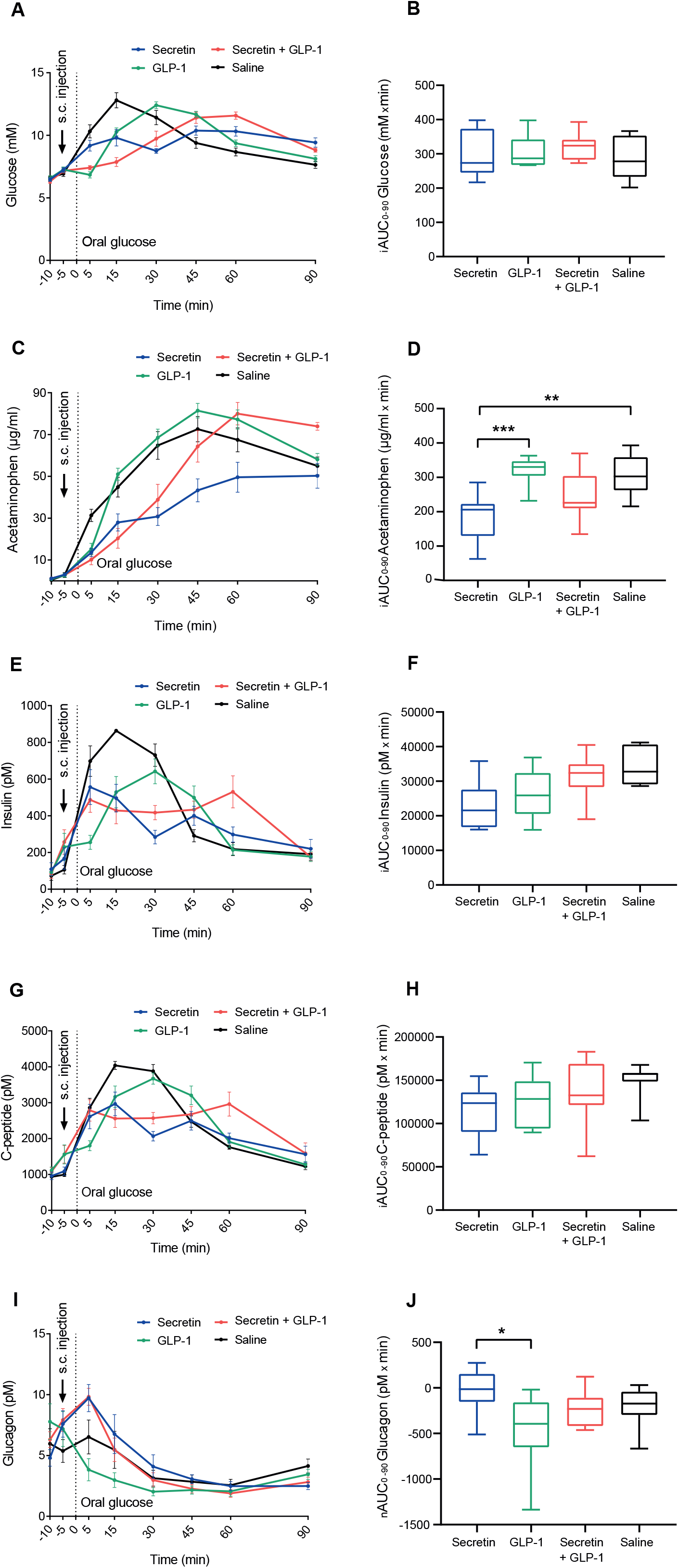
Effects of secretin and GLP-1 on glucose metabolism in non-sedated rats. Secretin (30 nmol/kg), GLP-1 (30 nmol/kg) or Secretin + GLP-1 (both 30 nmol/kg) were subcutaneously injected at timepoint −5 min. At timepoint 0, a mixture of oral glucose (2g/kg) and acetaminophen (25 mg/mL) was given. **A:** Blood glucose concentrations (mmol/L). **B:** Incremental area under the curve (iAUC_0-90_) of blood glucose concentrations shown in panel A. There were no significant differences in iAUC0-90 between the groups (P>0.77). **C:** Plasma acetaminophen concentrations (μg/mL). **D:** iAUC_0-90_ of acetaminophen concentrations. Acetaminophen concentrations in the group receiving secretin were significantly decreased compared to the control group (P<0.05) and to the group receiving GLP-1 (P<0.05). **E:** Plasma concentrations of insulin (pmol/L) **F:** iAUC_0-90_ of insulin concentrations. There were no significant differences in iAUC between the four groups (P>0.08). **G:** Plasma concentrations of C-peptide (pmol/L) **H:** iAUC_0-90_ of C-peptide concentrations. There were no significant differences between the groups (P>0.09). **I:** Plasma glucagon concentrations (pmol/L). **J:** iAUC_0-90_ of glucagon concentrations. Secretion of glucagon was significantly decreased in the group receiving GLP-1 compared to the group receiving secretin (P<0.05). There were no significant differences between all other groups (P>0.32). Data are shown as means ± SEM. P>0.05. *P<0.05, **P<0.01, ***P<0.001, by a One-way ANOVA with correction for multiple testing (Holm-Sidak), n=31 male Wistar rats (7-8 per group).

Plasma concentrations of insulin and C-peptide mirrored blood glucose levels (Figure 3E+G). However, in rats receiving both secretin and GLP-1, insulin and C-peptide peaked 5 minutes after the oral glucose gavage at which time point no change in blood glucose levels were observed (Figure 3A). When comparing iAUC_0-90min_ of insulin and C-peptide, there was a tendency towards a lowering effect of secretin and GLP-1, but neither reached statistical significance (Secretin vs. Saline; P_insulin_=0.07 and P_c-peptide_ =0.09, GLP-1 vs. Saline; P_insulin_ =0.09 and P_c-peptide_=0.55 and Secretin+GLP-1 vs Saline; P_insulin_=0.69 and P_c-peptide_=0.44) (Figure 3F+H).

Glucose administration lowered plasma glucagon concentrations in all groups (Figure 3I). However, secretin (either alone or in combination with GLP-1) treated rats had a significantly increased plasma concentration of glucagon during the first five minutes after administration (Secretin: 9.7±1.1 vs. Saline: 6.5±1.4 pmol/L, P=0.04 and Secretin+GLP-1: 9.8±0.7 vs. Saline: 6.5±1.4 pmol/L, P=0.03) but after 30 minutes, levels were similar to those in the control group. Glucagon n-AUCs were lower in the group receiving GLP-1 compared to saline group (P=0.08) and to the secretin-treated group (P<0.05) (Figure 3J).

### 3.6. Pancreatic delta cells express the secretin receptor (Sctr) and exogenous secretin increases somatostatin secretion from the perfused rat pancreas

To evaluate the potential metabolic effects of increased secretin release, we initially assessed tissue specific enrichment of the Sctr. We found that aside from the known expression of secretin receptors in the duodenum, pancreatic tissue contained the secretin receptor transcript (Figure 4A). However, since these values represent both expression in exocrine acinar/ductular tissue and islets, we next investigated if Sctr was present in human islets. Sctr was expressed in delta-cells and detectable at lower levels in alpha and beta-cells (Figure 4B). Sctr expression were compared to Glp-1r profiles, which, expectedly, showed high expression in the beta cells and, to a lesser extent, in the delta cells (Figure 4B). To investigate the translational relevance of rat islets to humans in regard to Sctr expression, we used an *in situ* hybridization approach. The Sctr was primarily found to be expressed in pancreatic delta-cells and to a lesser extent in the alpha and beta-cells of the rat (Figure 4C).

**Figure 4.**
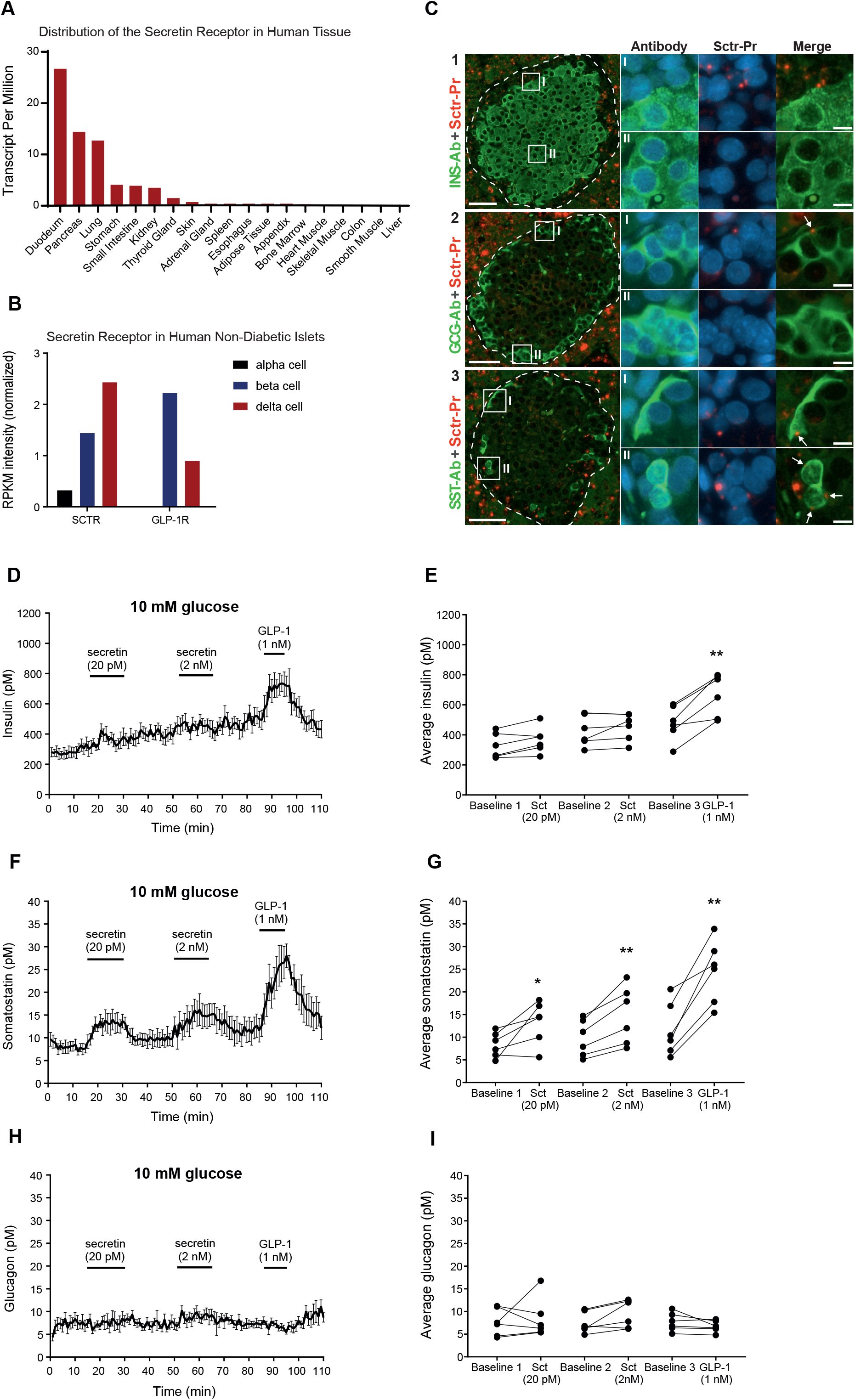
Expression of the secretin receptor (Sctr) and the effects of exogenous secretin on the perfused rat pancreas. **A: Expression profiles of Sctr in human tissues.** RNA-seq data from human tissues presented as Transcripts Per Million (TPM) (26). **B: Expression of Sctr and Glp-1r in human non-diabetic islets.** RNA-seq data from human alpha, beta and delta cells presented as Reads Per Kilobase Million (RPKM) (23–25). **C: Dual *in situ* hybridization and immunohistochemical staining of Sctr mRNA transcripts in rat pancreatic islets.** Representative images of pancreatic islets containing *in situ* hybridization stained Sctr mRNA transcripts (red) and immunohistochemically stained insulin (1), glucagon (2) and somatostatin (3) (green) cells with blow-up areas. Each red dot represents a single stained mRNA transcript. Arrows indicate overlap between Sctr mRNA staining and peptide hormone staining. Dashed line outline border between pancreatic islets and exocrine pancreas. Nuclei was visualized with DAPI counterstaining (blue). Arrows indicate staining overlap. Overview scale bar, 50 μm. Blow-up scale bar, 5 μm, n=4 male Wistar rats. **D-I: Effects of exogenous secretin on the perfused rat pancreas.** Secretin (20 pmol/L) was infused between minute 16-30 and Secretin (2 nmol/L) was infused between minute 51-65. GLP-1 (1 nmol/L) was infused between minute 86-95 as a positive control for insulin secretion **D:** Insulin concentrations (pmol/L). **E:** Mean concentrations of insulin at baseline (Baseline 1, 2 and 3) and following stimulation with either 20 pmol/L secretin, 2 nmol/L secretin or 1 nmol/L GLP-1. **F:** Somatostatin concentrations (pmol/L). **G:** Mean concentrations of somatostatin at baseline (Baseline 1, 2 and 3) and following stimulation with either 20 pmol/L secretin, 2 nmol/L secretin or 1 nmol/L GLP-1. **H:** Glucagon concentrations (pmol/L). **I:** Mean concentrations of glucagon at baseline (Baseline 1, 2 and 3) and following stimulation with either 20 pmol/L secretin, 2 nmol/L secretin or 1 nmol/L GLP-1. Data are shown as means ± SEM. P>0.05. *P<0.05, **P<0.01by a One-way ANOVA with correction for multiple testing (Holm-Sidak), n=6 male Wistar rats for all groups.

To examine whether secretin has an insulinotropic effect on the rat pancreas, we perfused the rat pancreas with 10 mmol/L glucose and administered secretin at a concentration matching post-RYGB concentrations as well as a supra-physiological concentration (20 pmol/L and 2 nmol/L respectively) (Figure 4D+E). GLP-1 was infused at the end of the experiment (1 nmol/L) as a positive control for insulin and somatostatin release (35–37). Secretion of insulin continuously increased throughout the experiment independent of secretin infusion (because of the elevated glucose concentration), and the baseline secretion of insulin was therefore calculated as the average secretion before and after secretin stimulation. Insulin did not increase in response to both of the applied concentrations of secretin (Baseline: 325±34 vs. Secretin (20 pmol/L): 365.9±35.2 pmol/L, P=0.13 and Baseline: 426.8±41.7 vs. Secretin (2 nmol/L): 454 ±36.8 pmol/L, P=0.56, *n=6*) (Figure 4E).

Consistent with the expression of the Sctr in rat and human delta-cells, secretin infusion significantly increased secretion of somatostatin (Baseline: 8.3±1.1 vs. Secretin (20 pmol/L): 13.3±1.9 pmol/L, P<0.05 and Baseline: 9.8±1.6 vs. Secretin (2 nmol/L): 14.9±2.6 pmol/L, P<0.05, *n=6*) (Figure 4F+G).

Glucagon secretion was low, due to the high glucose concentration in the perfusion buffer, and neither secretin nor GLP-1 infusions led to significant changes in its secretion (P>0.18, *n=6*) (Figure 4H+I).

## 4. Discussion

Here, we demonstrate that postprandial plasma concentrations of secretin are increased after RYGB surgery in obese individuals. This may result from the anatomical rearrangement of the small intestine which diverts luminal nutrients, like glucose, to more distal sites of the small intestine. Increased villus length after RYGB may also contribute the observed changes in plasma secretin concentrations (38). Tissue levels of secretin have been reported to be up-regulated in a rodent model of RYGB (39) but post-operative plasma concentrations of secretin have to our knowledge not been reported previously.

Furthermore, we show that secretin, besides in the duodenum, is expressed in the distal small intestine in humans and rats, and using a physiologically relevant model: the perfused rat intestine (31), we demonstrate that glucose is a strong stimulus for secretin release from the distal but not from the proximal rat small intestine. Using an inhibitor of SGLT1/2 activity (phloridzin) (34) we found that the molecular mechanism responsible for glucose-induced secretin secretion involves glucose absorption through the SGLT. SGLT-2 inhibition has gained major clinical interest due to its glucose lowering effect and reduced risk of cardiovascular disease (40). One may therefore speculate that patients treated with non-selective SGLT inhibitors would have an altered secretin secretion profile, similar to what has been reported for GLP-1 (41) but this remains to be explored. Previous attempts to show glucose-induced secretin secretion have been negative (7,42–44), probably because in these studies glucose was administered to the proximal rather than the distal part of the small intestine.

Glucose-stimulated secretin release from the distal small intestine is probably not of major physiological importance in un-operated humans since glucose mainly is absorbed in the proximal part of the small intestine. However, after intake of large carbohydrate rich meals, some intraluminal glucose may reach more distal parts of the small intestine (45), and secretion may under these circumstances play a role together with GLP-1 in improving glycemic control by inhibition of gastric emptying. This putative effect is supported by our in vivo data and consistent with previous findings (2–4).

Intraluminal acidification in the distal small intestine resulted in a larger secretory secretin response compared to the proximal intestine. The physiological relevance of this finding is not clear. The acidity of the gastric chyme entering the upper part of the small intestine is rapidly neutralized by a mixture of bile, mucosal - and pancreatic bicarbonate secretion, but the stimulatory effect of low pH in the distal small intestine on secretin-secreting cells may reflect a safety mechanism to reduce gastric emptying. The fact that also GLP-1 secretion was increased, may point to a more general effect, however, where hydrogen ions formed during digestion and perhaps fermentation in the mucosa micromilieu, stimulate ileal endocrine cells to activate the ileal brake. Further studies are required to investigate this new unexpected observation.

The observations regarding the effects of secretin on pancreatic islets are conflicting (6,7,46–50). Our data support that physiological levels of secretin regulate pancreatic secretion of somatostatin which is consistent with Sctr expression in rat and human delta-cells. Although Sctr was also expressed by beta-cells, secretin did not affect insulin secretion in the perfused rat pancreas whereas *in vivo* secretin actually led to lowering of plasma insulin levels early after an OGTT. The underlying reason for this awaits further investigation but this may be related to prolonged gastric emptying time and to an increased secretion of somatostatin which through paracrine effects may have overruled potential direct stimulatory effects of secretin on insulin secretion. Glucagon secretion was also not affected but this may be due to the already low levels when the pancreas was perfused in hyperglycemic conditions, thereby restricting the capability to detect further inhibition of alpha cells. However, to our surprise, there was a short-lasting increase in glucagon secretion after secretin injection *in vivo*, which was uninfluenced by the glucagonostatic effect of GLP-1 (51) and sympathetic stress (as this was not observed in the control group), as observed when similar doses of both peptides were injected. The underlying reason(s) is not clear.

## 5. Conclusion

Our study expands the current knowledge regarding secretin physiology and suggest that the RYGB-related increase in plasma secretin concentrations is mediated by nutrients reaching S-cells located in the distal small intestine. The physiological role of glucose-sensing S-cells warrant further studies but given secretin’s potential effects on islet secretion and whole-body metabolism (52) it may be speculated that secretin contribute to the metabolic effects of RYGB.

## Abbreviations

EECs: Enteroendocrine cells
GLP-1: Glucagon-like peptide 1
RPKM: Reads Per Kilobase Million
RYGB: Roux-en-Y gastric bypass
SGLTs: sodium-glucose co-transporters
TPM: Transcripts Per Million

## 6. Author contribution

Conceived and designed research: IMM, NJWA, and JJH. Performed experiments: DBA, KVG, REK, CBC, CØ, BH, and IMM. Provided clinical samples: CM, KNBM and SM. Provided Mass-spectrometry data: PL, RK, FR, and FG. Analyzed data: IMM and NJWA. Interpreted results of experiments: IMM and NJWA. Prepared figures: IMM and NJWA. Drafted manuscript: IMM, and NJWA. All authors edited and revised the manuscript and approved the final version.

## 8. Acknowledgement

The study was supported by a grant from the European Research Council (grant no. 695069) to JJH and by an Excellence Emerging Investigator Grant—Endocrinology and Metabolism (NNF19OC0055001) to NJWA.

