## Supplementary for "Secretin release after Roux-en-Y Gastric Bypass reveals a population of glucose-sensitive S-cells in distal small intestine"

[#]Shared co-last authors

**Corresponding authors:**

Professor Jens J. Holst, MD, DMSc, Department of Biomedical Sciences & Novo Nordisk Foundation Center for Basic Metabolic Research, Faculty of Health and Medical Sciences, University of Copenhagen, Blegdamsvej 3B, DK-2200 Copenhagen, Denmark; Telephone: +4535327518;

Or Assistant Professor Nicolai J. Wewer Albrechtsen, MD, PhD, Department of Clinical Biochemistry, Rigshospitalet & Novo Nordisk Foundation Center for Protein Research, Faculty of Health and Medical Sciences, University of Copenhagen. Blegdamsvej 9, DK-2100 Copenhagen, Denmark; Telephone +45 29649329;

### Supplementary Materials

##### *1. Development and evaluation of a secretin radioimmunoassay*

We obtained the following three secretin antisera: 1) ab103619 (Abcam, United Kingdom), 2) LS-C402778 (LSBio, Seattle, USA), 3) A C-terminally directed secretin antibody (codename 5595-3) kindly provided by Professor Jan Fahrenkrug (Department of Clinical Biochemistry, Bispebjerg Hospital, Copenhagen, Denmark). The binding region of antibody 1 and 2 was not known.

Cross-reactivity in the secretin assay was assessed for the following peptides: Glucagon 1-29 (Cat. No.: H-6790, Bachem), oxyntomodulin (Cat. No.: H-6058, Bachem), and glicentin (GenScript, Piscataway, USA, custom made service no. SC1208) as well as CCK-8 (Cat. No.:4033010, Bachem), recombinant human insulin (Cat. No.: ab86658, Abcam), neurotensin (Cat. No.: N6383, Sigma) and human PYY 3-36 (Cat. No.:4018880, Bachem).

Commercially available secretin ELISAs (LS-F27215 (LSBio) and CEB075Hu (Cloud Clone, Texas, USA) were included in our assay evaluation for comparison with our in-house developed assay. Assays were evaluated according to the CLSI (Clinical and Laboratory Standards Institute) guidelines for Immunoassay (I/LA23-A, I/LA21-A2, and EP24-A) as previously described (2).

Recovery in percent was calculated as the measured concentration in the individual, spiked plasma sample minus the concentration in the corresponding non-spiked plasma sample divided by the theoretical spiked concentration of peptide and multiplied by 100.

Specificity was analyzed from recovery experiments. Known amounts of each peptide (5–300 pmol/L) were added to separate aliquots of healthy human plasma (a reserve plasma pool from a previous study (1) was used). One aliquot was measured in duplicate, using two separate assay runs or kits for each of the two commercial ELISAs.

Sensitivity was estimated by determining the lowest concentrations of added peptide that could be measured as being significantly different from zero (by paired analysis of three duplicate determinations) using a plasma pool spiked with 0, 1, 2, 5 pmol/L of human secretin. Precision was determined from multiple measurements of identical samples.

For all kits, the manufacturers' instructions were followed closely, including recommendations for sample preparation by extraction (purification columns and buffers).

***2. Peptide extraction in rat tissue biopsies***

The biopsies were cleaned in PBS and snap frozen on dry ice before storage at -80°C. Peptides were extracted using 1% trifluoroacetic acid (cat no. TS-28904, Thermo Fisher Scientific, USA) and homogenized with a bead mill (TissueLyzer, Qiagen Instruments AG, Switzerland) and a 5 mm steel bead. Peptides were purified using tc18 cartridges (cat no. 036810 Waters, USA). Before analysis of secretin and GLP-1 peptide content, samples were reconstituted in 1 ml assay buffer (80 mmol/L phosphate buffer, 10 mmol/L EDTA, 0.6 mM Thimerosal (cat no. T-5125, Sigma Aldrich, Denmark), 0.1% Human Serum Albumin (cat no. 12666, Merck KgaA, Germany), pH 7.5). Peptide content was normalized to tissue weight (pmol/g).

***3. Immunohistochemistry***

Tissue were paraffin-embedded on a HistoStar embedding workstation (Thermo Fisher Scientific, Massachusetts, USA), cut on a HM355S automatic microtome (Thermo Fisher Scientific, Massachusetts, USA) (4 μm) and mounted on cover slides. Next, the slides were heated at 60°C for 60 minutes to remove the paraffin and placed in a TissueClear bath for 10 min followed by several baths at decreasing ethanol concentrations, from 99-70%, to hydrate the samples. Heat-induced antigen retrieval was done by microwaving samples for 15 min, in a T-EG buffer (pH 6, 0.5mmol/L Trisma base, Cat. no. T6066, Sigma-Aldrich, Missouri, USA, 1mM EGTA, Cat. no. E4378, Sigma-Aldrich, Missouri, USA) and leaving them in the warm buffer for 20 min. Samples were washed with PBS, and blocked in 2% (w/v) BSA for 10 min. Then incubated with the primary antibody (1:20000, Polyclonal Rabbit Anti-Secretin (5585-3), a generous gift from J. Fahrenkrug at Bispebjerg Hospital, Copenhagen, Denmark) for 1 hour, washed three times 1 minute in wash buffer (EnV FLEX wash buffer 20x, Cat. no K800721-2, Agilent, Glostrup, Denmark) before incubating samples with secondary antibody (1:200, Polyclonal, biotinylated Anti-rabbit, Cat. no. BA-1000, Vectorlabs, California). Samples were washed three times in wash buffer, incubated in 3% (v/v) $H_2O_2$ for 8 min at 37°C, followed by three washes, and incubation with the Vectastain Elite ABC complex (mixed 30 min prior to use according to supplier instructions, Cat. no. PK-6100 Vectorlabs, California, USA). Another three washes were carried out before developing the sections by incubation with DAB+ (mixed according to supplier instructions, Cat no. SK4105, Vectorlabs, California, USA), for 15 min. Sections were washed in Milli Q water for 5 min and bathed in hematoxylin for ~30 seconds. The slides were rinsed in a Milli Q water bath to remove any excess hematoxylin before mounting coverslips with PERTEX mounting medium (HistoLab, Västra Frölunda, Sweden).

##### *4. Isolated perfused rat small intestine and pancreas*

Non-fasted rats were anaesthetized with a subcutaneous injection of Hypnorm/Midazolam. The proximal and the distal small intestines (45±5 cm) were isolated by ligating the blood supply and removing the colon and either the proximal or the distal half of the small intestine. A catheter was inserted into the superior mesenteric artery and the intestine was perfused at a constant flow rate of 7.5 mL/min with a heated (37°C), oxygenated (95% $O_2$, 5% $CO_2$) perfusion buffer (pH 7.4). To remove intestinal contents, a plastic tube was placed in the lumen of the intestine and the intestine was gently flushed with pre-warmed saline (0.9%). Throughout the experimental protocol, a constant luminal flow of heated saline was applied via a syringe pump (0.25 mL/min). The venous effluent was collected from a catheter inserted into vena portae.

Pancreas was isolated by removing the large and small intestine, the spleen and the stomach after ligation of their specific vessels. The renal arteries were ligated and the abdominal aorta was ligated below the diaphragm. A catheter was inserted into the abdominal aorta proximal to the renal arteries and the pancreas was perfused at a constant flow rate of 5 mL/min. The venous effluent was collected from a catheter in vena portae. As soon as proper flow was apparent, the rats were euthanized by perforation of the diaphragm. For stabilization, the organs were perfused for 25 minutes before initiation of the experimental protocol.

Perfusion buffer was a Krebs-Ringer bicarbonate buffer supplemented with 5% (w/v) dextran T-70 (Pharmacosmos, Denmark), 0.1% (w/v) bovine serum albumin (cat. no. 1.12018.0500, Merck, Denmark), 5 mmol/L of each pyruvate, fumarate and glutamate (Sigma Aldrich, Denmark). The perfusion buffer contained 3.5 mmol/L glucose in the perfused intestine and 10 mmol/L glucose in the perfused pancreas.

The venous effluent was collected for 1 min periods via the draining catheter using a fraction collector. The samples were immediately put on ice and stored at -20°C until analysis.

##### *5. In situ hybridization*

Pancreases were excised from non-fasted rats and transferred to fresh 4% paraformaldehyde (PFA), PBS for 24h at room temperature. Tissue was then transferred to 70% ethanol followed by infiltration (Shandon Excelsior, Thermo Fisher), and embedded in paraffin blocks. Sections of 4 μm were cut using a microtome (RM2125, Leica) and mounted on slides (superfrost Plus Slides, Thermo Scientific). The slides were baked at 60 °C for 1h and placed in xylene and ethanol for dewaxing. The *in situ* hybridization was performed using mRNA specific probes (Supplementary Table 3) following the manufacturers' manual of a commercially available *in situ* hybridization kit, RNAscope 2.5 HD detection (Cat.no. 322360, Advanced Cell Diagnostics, Milan, Italy) in combination with immunofluorescence. After the *in situ* hybridization procedure, the slides were incubated with blocking buffer (2 % BSA, PBS for 10 minutes) followed by incubation with primary insulin, glucagon or somatostatin antibody in blocking buffer (Supplementary Table 2) over night at 4 °C. The day after the slides were washed and incubated with Alexafluor 488 conjugated secondary antibodies for 1 hour at room temperature (Supplementary Table 2). Coverslips were then mounted with ProLong Gold 32 Antifade Mountant with DAPI (P-3693 I, Thermofisher). The slides were analyzed using a Zeiss Widefield fluorescence microscope and AxioCam 506 mono camera or a Zeiss Axio Scan.Z1 Slide Scanner and AxioCam MRm camera.

### Supplementary Figures

**Supplementary Figure 1:** Evaluation of an in-house secretin assay

In-house developed assay calibrator curve showing 95% CI and corresponding details of the non-linear regression fit (a 4-parametric logistic regression model) below.

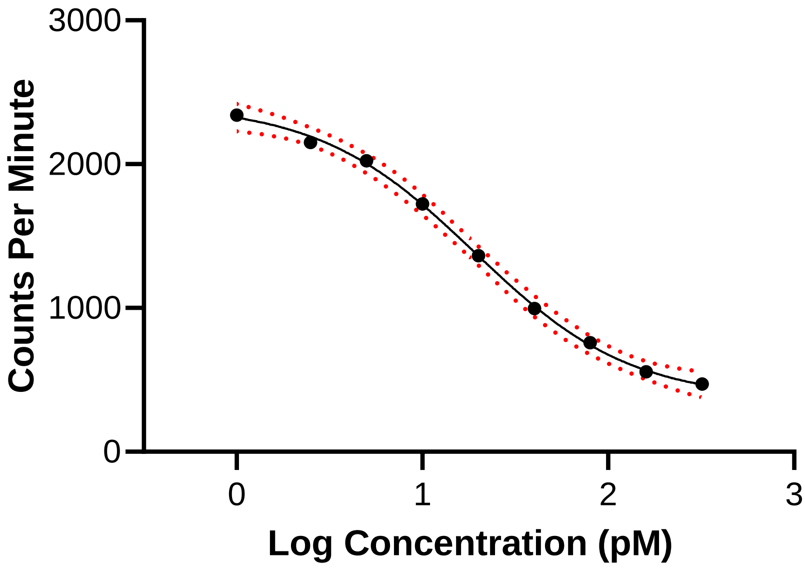

| | |
|---|---|
| Best-fit values | |
| Top | 2417 |
| Bottom | 361,6 |
| LogIC50 | 1,279 |
| HillSlope | -1,033 |
| IC50 | 19,01 |
| Span | 2056 |
| 95% CI (profile likelihood) | |
| Top | 2271 to 2651 |
| Bottom | 118,7 to 507,6 |
| LogIC50 | 1,172 to 1,387 |
| HillSlope | -1,354 to -0,7423 |
| IC50 | 14,87 to 24,40 |
| <u>Goodness of Fit</u> | |
| Degrees of Freedom | 50 |
| **R squared** | **0,9645** |
| Sum of Squares | 903621 |
| Sy.x | 134,4 |

**Supplementary Figure 2:** Bland-Altman plots of assay B versus assay A, and assay C versus assay A

Assay A: In-house RIA, Assay B: LS-F27215 (LSBio) and Assay C: CEB075Hu (Cloud Clone, Texas, USA). Human secretin was spiked in human plasma at different final concentrations and a bland-Altman analysis was performed using GraphPad Prism. The differences (%) was visualized as shown below for the two comparisons to the in-house developed assay.

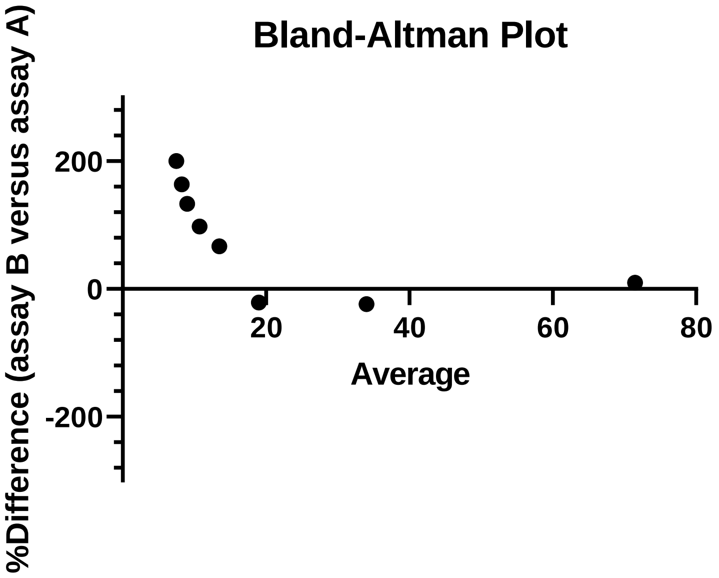

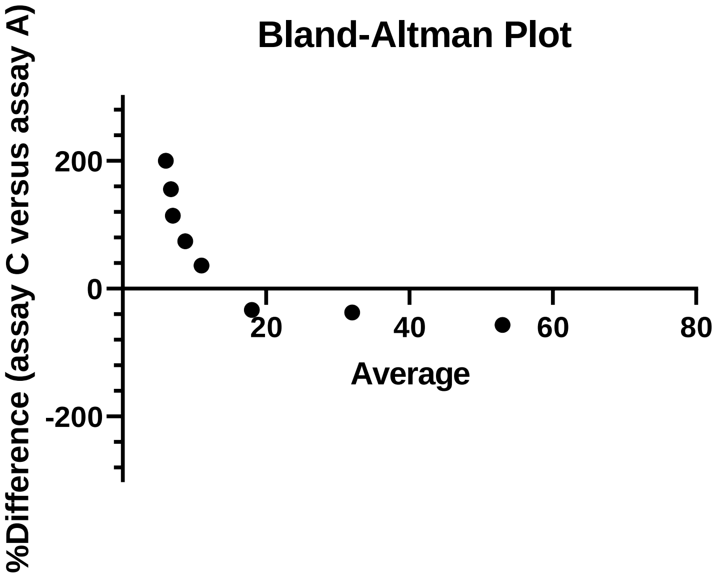

### Supplementary Tables

**Supplementary Table 1:** Anatomical Definitions

| Nomenclature used in the manuscript | Anatomical Location |
|---|---|
| Esophagus | 0.5 cm proximal to esophageal sphincter |
| Ventricle | 1 cm proximal to pyloric sphincter (antrum) |
| Duodenum | 1 cm distal to pyloric sphincter |
| Proximal Jejunum | 6 cm distal to pyloric sphincter |
| Distal Ileum | 1 cm proximal to ileocecal junction |
| Colon | Mid part of colon |

**Supplementary Table 2**: Antibody Table

| Target protein | Manufacturer/ Provider and cat. no. | Host | IHC dilutions for *in situ* hybridization | IHC dilutions for secretin localization |
|---|---|---|---|---|
| **Insulin** | In-house, 2006-4 | Guinea Pig | 1:3000 | - |
| **Somatostatin** | In-house, 1759 | Rabbit | 1:3000 | - |
| **Glucagon** | Abnova, MAB1500 | Mouse | 1:300 | - |
| **Secretin** | A gift from Fahrenkrug,5585-3 | Rabbit | - | 1:20000 |
| **CF488A anti-Guinea pig IgG** | Biotium, 20169 | Donkey | 1:200 | - |
| **Alexa fluor 488 anti-rabbit IgG** | Abcam, ab150109 | Donkey | 1:200 | - |
| **Alexa fluor 488 anti-rabbit IgG** | Abcam, ab150081 | Goat | 1:200 | - |
| **Biotinylated anti-rabbit IgG** | Vector Laboratories, BA-1000 | Goat | - | 1:200 |

**Supplementary Table 3:** Probe Table

| Target mRNA | Target species | Manufacturer/ Provider and cat. no. |
|---|---|---|
| **Secretin receptor (Sctr)** | Rattus norvegicus | ACD biotechne, 580701 |
| **Peptidylprolyl isomerase B (Ppib)** | Rattus norvegicus | ACD biotechne, 313921 |
| **Dihydrodipicolinate reductase (DapB)** | Bacillus subtilis strain SMY | ACD biotechne, 310043 |
